# The non-random assembly of functional motifs in plant-pollinator networks

**DOI:** 10.1101/2022.04.06.486621

**Authors:** Jose B. Lanuza, Alfonso Allen-Perkins, Ignasi Bartomeus

**Affiliations:** School of Environmental and Rural Science, University of New England, Armidale, New South Wales 2350, Australia; Estación Biológica de Doñana (EBD-CSIC), E-41092 Seville, Spain; Departamento de Ingeniería Eléctrica, Electrónica, Automática y Física Aplicada, ETSIDI, Universidad Politécnica de Madrid, 28040 Madrid, Spain

**Keywords:** plant-pollinator networks, plant-pollinator interactions, network motifs, indirect interactions, plant traits, floral visitors

## Abstract

Ecological processes leave distinct structural imprints on the species interactions that shape the topology of mutualistic networks. Detecting those relationships is not trivial since they go beyond pairwise interactions, but may get blurred when considering full network descriptors. Recent work has shown that the network meso-scale can capture this important information. The meso-scale describes network subgraphs representing patterns of interactions between a small number of species (i.e., motifs) that constitute the building blocks of the whole network. Despite the possible implications of network motifs to better capture species interactions, they remain overlooked in natural plant-pollinator networks. By exploring 60 empirical plant-pollinator networks from 18 different studies with wide geographical coverage we show that some motifs are consistently under- or over-represented worldwide, suggesting that the building blocks of plant-pollinator networks are not random. Furthermore, we find that distinct motif positions describing species ecological roles (e.g., generalisation and number of indirect interactions) are occupied by different plant and floral visitor groups on both trophic levels. Bees appear less frequently in specialised motif positions with high number of indirect interactions, while the rest of floral visitor groups are infrequent in generalised motif positions with low number of indirect interactions. All plant groups tend to be over-represented on specialised motif positions, except tall plant species with separated sexes (e.g., woody dioecious or monoecious species), which are more frequent on motif positions with low number of indirect interactions. Interestingly, the realized combinations of different species groups within a motif can not be retrieved from their joint probability distributions, indicating that group combinations are not random either. Our result highlights the non-random structure of the meso-scale on plant-pollinators networks and the association of different plant and floral visitor groups with certain motifs that involve different ecological roles at a macro-ecological scale.

## Introduction

Ecological communities are formed by a plethora of interacting species that form networks of interactions. Because of the tremendous complexity of these ecological networks, species interactions are generally condensed in metrics that summarize this information (Guimarães, 2020). Plant-pollinator interactions are no exception, and they are often studied with indices that aggregate the information at the network (macro-scale) and species level (micro-scale). For example, plant-pollinator network level approaches have identified common invariant structural properties across networks, including a degree distribution that decays as a power law (Jordano, 1987), nestedness (Bascompte et al., 2003), or modularity (Olesen et al., 2007). In addition, the local species position within the network can define its degree of specialisation (Blüthgen et al., 2006) or its role in connecting the rest of the community (Olesen et al., 2007). Despite the unquestionable progress with the use of these metrics at both network and species level, condensing complex information into a single metric implies the loss of relevant ecological information that obscures the understanding of species interactions (Cirtwill et al., 2018; Simmons, Cirtwill, et al., 2019).

Traditionally, plant-pollinator research has focused on direct interactions but overlooked indirect interactions (i.e., the mediated effect between two species by a third species), such as facilitative or competitive interactions between plants for pollinators (Carvalheiro et al., 2014; Moeller, 2004; Sargent & Ackerly, 2008). Despite the widespread nature of indirect interactions in ecological communities (Strauss, 1991), plant-pollinator research often fails to finely capture those indirect interactions with the conventional analytical tools that condense the information either by species (e.g., degree) or in global topological indices (e.g., nestedness). Nonetheless, the emerging framework of network motifs (small sub-networks observed within a given network which are often referred as its building blocks; Milo et al., 2002) allows to consider both direct and indirect interactions (Simmons, Cirtwill, et al., 2019). The analysis of network motifs in plant-pollinator networks have revealed that the different ecological processes that govern species interactions (e.g., species abundances versus trait-matching) can lead to different patterns of indirect interactions (Simmons et al., 2020). Yet, the global patterns of indirect interactions in real plant-pollinator networks are still unknown (e.g., over- and under-represented motifs).

By linking the structural properties of the meso-scale with the species’ ecology we can advance in our understanding on the determinants of species interactions. For instance, different motifs can have different ecological meanings (Simmons, Cirtwill, et al., 2019) and the position within a motif can determine the species’ functional role (Baker et al., 2015; Stouffer et al., 2012). Yet, our understanding of the ecological implications of network motifs in real plant-pollinator networks remains in its infancy. Moreover, it is unclear how the species ecology and life-history traits determine its functional role within the network of interactions (Coux et al., 2016). For example, large pollinators can forage larger distances (Greenleaf et al., 2007), deposit greater pollen quantities (Földesi et al., 2021) and handle complex zygomorphic flowers in comparison with small pollinators that are restricted to lower floral complexity (Gong & Huang, 2009). Despite these obvious differences in pollinator behaviour, how contrasting species’ ecology translates into different interaction patterns is still unknown. Similarly, recent empirical findings indicate that the meso-scale is the best descriptor of plant reproductive success (Allen-Perkins et al., 2021), but little is known about how plants reproductive strategies shape their position within the network of interactions. Although some studies have evaluated plant reproductive strategies in plant-pollinator networks (Lázaro et al., 2020; Tur et al., 2013), this is an often overlooked aspect in a community context (Devaux et al., 2014) and rarely incorporated into plant-pollinator network studies. Hence, exploring how the main plant and floral visitor life-history strategies integrate with the emergent framework of network motifs could help to progress knowledge on the ecological processes that shape plant-pollinator interactions.

Here, we used 60 empirical networks from 18 different studies and 14 countries to evaluate the structure of the meso-scale in plant-pollinator networks. From these networks we extracted plant and floral visitors species and determined the principal groups that define their main life-history strategies. To obtain plant groups, we used a comprehensive dataset that included floral, reproductive and vegetative traits compiled in a larger set of plant species. Floral visitor groups were divided by the main taxonomic groups that differed in life form and behaviour. Once we split the different plant-pollinator networks into their motif elements, we explored: (i) if there is a common invariant structural property in the overall motif networks (i.e., over- and under-represented motifs); (ii) which plant and floral visitor groups are over- and under-represented in different motif positions associated with specific ecological roles; and, (iii) if there are over- and under-represented plant and floral visitor group combinations within a motif.

## Methods

### Overview

To investigate the structure of the meso-scale in plant-pollinator networks, we gathered published plantpollinator networks with worldwide distribution. Then, we classified the different plant and floral visitor species from these networks into groups that summarise their main life-history strategies. After, we calculated the normalised motif frequencies and investigated if these frequencies are similar or not to the ones expected by chance with the help of null models. Then, for each group of plants and floral visitors we investigated the ecological implications of their overall representation on the different motif positions, by comparing, on the one hand, the observed (normalised) frequencies of each group in the different motif positions with the frequencies of their simulated counterparts, and, on the other hand, the differences among the patterns of connections of motif positions. Finally, we compared if there are differences in the observed frequencies of motif combinations from the empirical networks and the expected frequencies of motif combinations calculated as the product of the probabilities of the different plant and floral visitor groups within the motif.

### Plant-pollinator studies

We have compiled 60 plant-pollinator networks from 18 different studies (**Supporting Information Table S1**). All studies sampled plant-pollinator interactions in natural systems and were selected based on wide geographical coverage (**Supporting Information Figure S1**) and presence of interaction frequency as a measure of interaction strength. In total, there were 503 plant species, 1,111 floral visitors species and 6,248 pairwise interactions registered. For ease of data manipulation, plant and floral visitor species names were standardize with the help of the R package *taxize* version *0.9.99* (Chamberlain et al., 2020). All analyses and data manipulation were conducted in R *version 4.0.5* (R Core Team, 2021).

### Plant and floral visitor groups

First, plant species were grouped through hierarchical clustering into the optimal number of functional groups that summarized the main plant reproductive strategies. For this, we used the trait dataset collated in Lanuza et al. (2021) that comprised a total of 1506 species including the 503 species considered in this study (**Supporting Information Table S2**). This dataset consisted on 8 floral, 4 reproductive and 3 vegetative traits excluding traits with high percentage of missing values (over 30%; **Supporting Information Table S3**). We opted to calculate the plant functional groups on this larger set of species because of the higher accuracy when delimiting functional groups with that many variables and species (Dolnicar et al., 2014). To feed the clustering analysis, we calculated the distance between the different qualitative and scaled quantitative variables with Gower distance (Gower, 1971). For this, we used the function *gowdis* with method *ward.D2* from the *FD* package version *1.0-12* (Laliberté et al., 2014). Finally, we conducted hierarchical clustering with the function *hclust* from the *R* stats package version *4.0.5* and calculated the optimal number of clusters with the function *kgs* from the *maptree* package version *1.4-7* (White & Gramacy, 2009).

Second, floral visitors were aggregated into taxonomic groups based on taxonomic rank as done similarly in other plant-pollinator studies (Fenster et al., 2004; Ollerton et al., 2009). We opted to divide floral visitors on the taxonomic rank level and not with functional traits because the main orders of floral visitors differed in form and behaviour and had lower superior taxonomic complexity (i.e., floral visitors had 6 orders versus plants that had 38). Thus, this allowed us to group floral visitors into groups that represented approximately the main life strategies of the possible pollinators: (i) bees (Hymenoptera-Anthophila), (ii) non-bee Hymenoptera (Hymenoptera-non-Anthophila), (iii) syrphids (Syrphidae-Diptera), (iv) non-syrphids-Diptera, (v) Lepidoptera and (vi) Coleoptera. However, a minor set of species belonged to other taxonomic groups that were considered in analyses but not discussed further because of their low representation in the full set of networks (3.55% of the total interactions recorded). These taxonomic groups were ‘lizards,’ ‘birds’ and ‘other insects.’ This last group was formed by a mix of uncommon insect taxa on the full set of networks.

### Overall motif patterns

Following previous work (Simmons, Cirtwill, et al., 2019; Simmons et al., 2020), we broke down the plantpollinator networks into their constituent motifs that can capture both direct and indirect interactions. Prior to analyses, we turned the quantitative networks into qualitative (or binary) ones, where interactions are present or absent. This approach was preferred since quantitative motif analysis have been developed just recently and more work is needed to fully understand and interpret the information that they can convey (Simmons et al., 2020). All analyses were run considering singletons (species with only one interaction detected) but we conducted a second exploration without singletons (64.98% of interactions) to evaluate the effect of rare species on the observed patterns. Results without singletons are qualitatively consistent and are not further discussed in the main text (**Supporting Information Figures S2 and S3**).

We calculated the frequency of all motifs up to five nodes (17 different motifs in total; see **Figure 1**) for each empirical network, by using the *bmotif* package version *2.0.2* (Simmons, Sweering, et al., 2019). We focused exclusively on motifs up to five nodes in our analyses for computational reasons and for the ease of visualization and interpretation of a reduced set of less complex network motifs. Throughout the entire manuscript, floral visitors occupy the top nodes of the different motifs and plants the bottom ones. To control for variation in network size, motif frequencies were normalised as a proportion of the total number of motifs within each motif class (i.e., 2-species motifs, 3-species motifs, 4-species motifs and 5-species motifs). In addition, we excluded 2-species (or nodes) motifs from our analyses because its normalised frequencies would always equal one. Note that the aim is not comparing different networks, which may differ in sampling effort, but correctly describing the structure of each individual network. Like Simmons et al. (2020), we classified the different motifs by their average path length (mean number of links between all pairs of nodes) into four different groups: (i) complete or strong 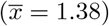, (ii) fan or medium-strong 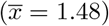, (iii) assymmetric complete or medium-weak 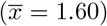 and (iv) core-peripheral or weak 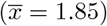. In general terms, plant and floral visitor groups on more densely connected motifs (e.g., complete or fan) will experience lower number of indirect interactions in contrast to groups on less densely connected ones (e.g., medium-weak or weak).

**Figure 1.**
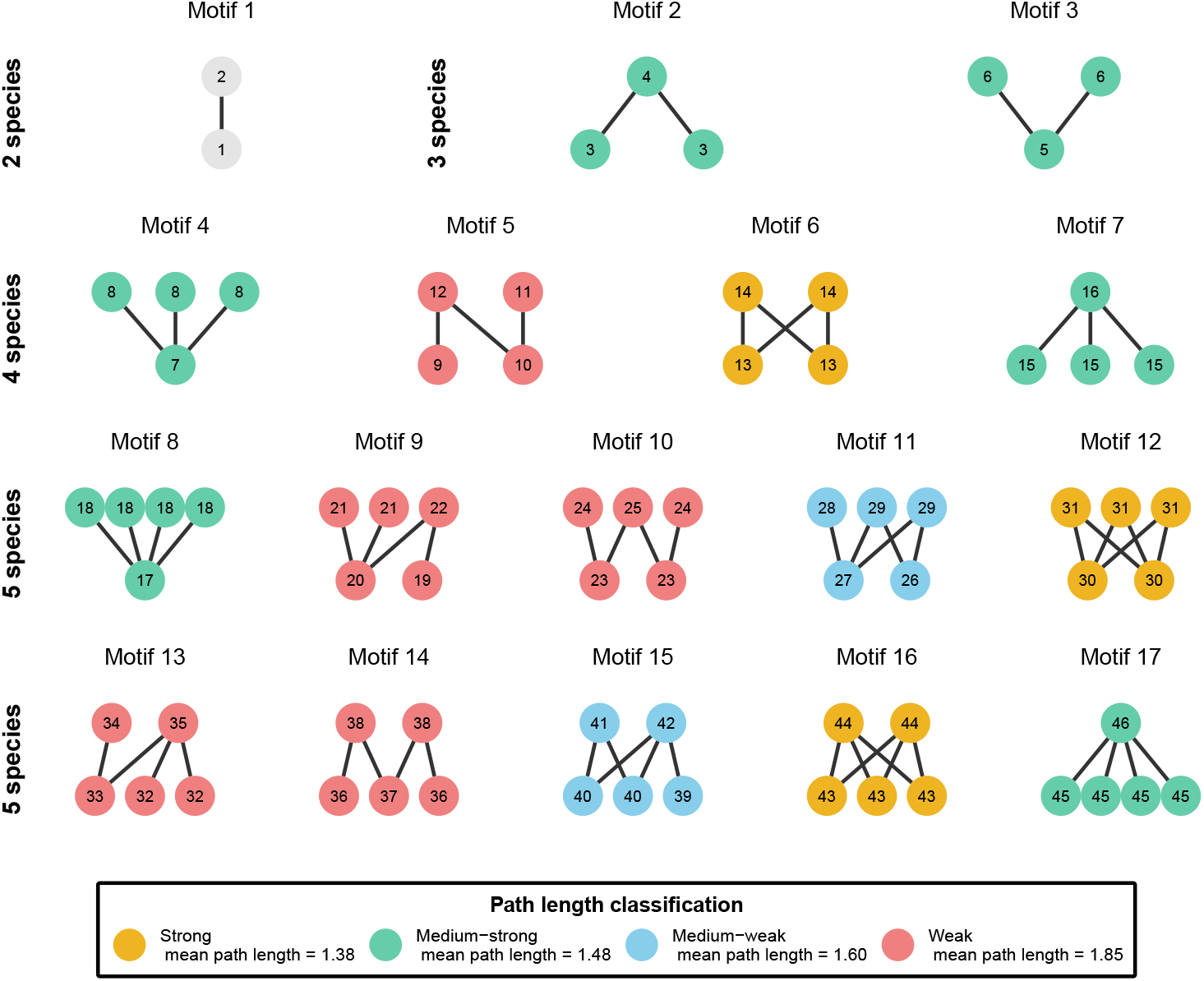
Adapted figure of Simmons et al., (2019). All possible motif network structures from two to five species in bipartite networks grouped by the different motif classes (2-species, 3-species, 4-species and 5-species). There is a total of 17 possible motifs with 46 different positions denoted within each node. The different motifs are aggregated by their average path length classification wich goes from densely connected motifs (strong motifs coloured in yellow) to less connected motifs (weak motifs coloured in red). Note that two node motifs are excluded from this classification.

To assess the significance of the observed frequencies, we created 1,000 simulated networks for each binary network using the *nullmodel* function and the *vaznull* model in the *bipartite* package version *2.14* (Dormann et al., 2009). Generated networks had the same number of plants and floral visitors, as well as the same connectance of their corresponding empirical networks. We discarded null models with more constraints (e.g., constraining for species degree) because the permutations did not generate sufficient variablity and provided highly similar or identical networks to the original ones. After extracting the motif frequencies from the simulated networks, for each motif type and empirical network, we calculated the percentage of simulated networks whose frequencies were smaller than the ones observed, that is, we estimated the percentile of the observed motif frequencies. Motifs whose percentile is close to 0 or 100 are under- or over-represented in the empirical networks, respectively, and thus they cannot be predicted by connectance and the number of species alone. To summarize general patterns across networks, we used an intercept-only linear mixed model (LMM) per motif with the help of the *lme4* package version *1.1-21* (Bates et al., 2015), where the response variable was the observed motif percentile per network. Note that as we are only interested in modelling the intercept value, no predictors were specified. In these models, we used the study identifiers in **Supporting Information Table S1** as a random intercept. By doing so, we obtained estimates of the average motif frequency, but controlling for the variation at the study level.

### Over- and under-represented groups on motif positions

We calculated which plant and floral visitor groups were over- or under-represented in the different motif positions by comparing position frequencies of empirical networks with those of their corresponding simulated counterparts. To estimate the position frequencies of each plant and floral visitor group in a given network, we added the frequencies of those species that belong to the same group, and then, we normalised the resulting frequencies by dividing them by the total number of times that a corresponding group appears in any position within the same motif size class (i.e., 2, 3, 4 or 5 species motifs). Then, we calculated the percentile of the observed position frequencies for each group and network, just like we did with motif frequencies. To outline the general patterns of position frequencies across networks and groups, we fit a LMM per motif position, where the response variable was the observed position percentile per network. We used the group identifier as an explanatory variable and the study identifiers as a random intercept. Hence, we assessed the average motif frequency per species group, after controlling for the variation at the study level. Finally, we visualized with the help of the *ComplexHeatmap* package version *2.6.2* (Gu et al., 2016) over- and under-representation of plant and floral visitor groups on the different motif positions.

To understand if the different plant and floral visitor groups are associated with motif positions that represent different network roles, we characterized the roles of each motif positions for plant and floral visitor groups in terms of their specialisation and number of indirect interactions. To that end, we calculated the specificity of each motif position following Poisot et al. (2015) as *s_i_* = *L* – *l_i_/L* – 1, where *L* is the total number of groups and *l_i_* the number of interaction partners of the group *i* in a given motif. Values of the vector *s_i_*, range between 1, complete specialisation (1 partner within the motif) and 0 complete generalism (all possible partners within the motif). Then, because all nodes within a motif are connected, we calculated the number of indirect interactions (*n_i_*) for each motif position and trophic guild (i.e., plants or floral visitors) as *n_i_* = *z_i_* – 1, where *z_i_* is the total number of nodes for a trophic guild in a given motif. Thus, we considered exclusively indirect interactions within plants though shared floral visitors or within floral visitors though shared plants. Finally, we analysed in a single model how specialization and the number of indirect interactions were associated with the observed percentiles for each trophic guild and motif position (see paragraph above). For this, we fitted a LMM model with the help again of the *lme4* package where we considered specialization and number of indirect interactions as response variables with the interaction by plant and floral visitor group for both response variables. Further, we considered as random factor motif position nested within study system. All model diagnostics were conducted with the help of the *Dharma* package (Hartig & Hartig, 2017) version *0.3.3.0* and model outputs were visualized with the *visreg* package (Breheny et al., 2020) version 2.7.0.

### Over- and under-represented group combinations of motifs

Finally, we studied which combinations of plant and floral visitor groups tend to appear together within the same motifs (up to five nodes). That is, which combinations are over- or under-represented. This analysis used 57 out of the 60 networks available due to computational limitations to identify all the nodes in the motifs of the three networks with the highest number of links. To do so, for each of the 53,250 possible motif combinations, we estimated the observed and the expected probability of finding those combinations in empirical networks, respectively. Then, we determined whether the observed probabilities are likely to come from the expected probabilities or not. To calculate the observed probability 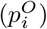 of the different groups within a motif *i* (e.g., motif 3, ‘bee’ + ‘bee’ + ‘selfing herbs’), we divided the number of times that *i* appears in our set of empirical networks 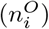 by the sum of the number of times that each possible combination appeared: 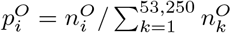. To estimate the expected probability 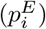 of a given motif combination *i*, we assumed that the probability of finding a given group *x* in the position *α* is independent of the probabilities associated with other positions and groups in that motif. Therefore, the expected probability 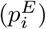 of the motif combination *i* is given by the product of the probabilities of its constituents 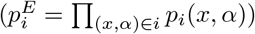. Then, by controlling for the variation at the network level with an LMM (see below), we calculated the average number of times that the group *x* appears in the position *α* (denoted as *n_i_*(*x, α*)). To obtain the expected probability *p_i_*(*x, α*) we divided the number of times that the group *x* appears in the position *α* by the the total number of times that any group was observed in that position (that is, *p_i_*(*x, α*) = *n_i_*(*x, α*)/∑_*k*_ *n_i_*(*k, α*)). The average number of times that the group *x* appears in the position *α*(*n_i_*(*x, α*)) was calculated with a LMM for each position, where the response variable was the number of times that a group was observed at a given position in each real-world network, the explanatory variable was the group identifier, and the random intercept was given by network identifiers nested within the study identifiers.

Once we obtained 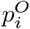 and 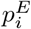, we used a simulation approach to determine whether the former is likely to come from the latter or not. This approach was preferred since the large number of possible motif combinations and the small probabilities for some of them advise against using an exact test of goodness-of-fit or a Chi-square one. Specifically, we created 1,000 random samples with repetition of possible motif combinations, where each sample contained 10 million elements and, for each combination, the probability of being selected was equal to its expected probability. From those random samples, we extracted the mean and the standard deviation of the expected probability of *i*, denoted as 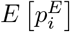 and 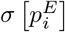, respectively, and calculated the z-scores of 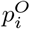 as 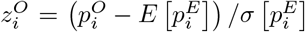, for those motif combinations with 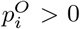. According to the usual interpretation of z-scores, combinations with 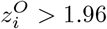 are over-represented, whereas those with 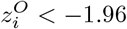 are under-represented, at the 95% confidence level. Notice that we focused on combinations with 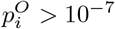 because we do not have enough numerical resolution to accurately detect whether combinations that took place only once are under-represented (due to the limited size of our random samples).

## RESULTS

### Plant and floral visitor groups

The hierarchical cluster analysis divided the dataset into five different groups with different and overlapping characteristics (**Supporting Information Figure S4** and **Figure S5**). The first group referred to as ‘selfing herbs’ consisted on herbs with hermaphroditic flowers with high levels of autonomous selfing. The second group named ‘small outcrossing perennials’ had small perennial species with a mix of life forms (i.e., trees shrubs and herbs) with outcrossing hermaphroditic flowers. The third group referred to as ‘self-incompatible perennials with large flowers’ comprised perennial species with a mixed of life forms and large self-incompatible hermaphroditic flowers with high number of ovules. The fourth group named ‘tall plants with small unisexual flowers’ had the tallest species, highest proportion of shrub and tree life forms, dioecious and monoecious breeding systems, small flowers and the highest numbers of flowers and inflorescences per plant. Finally, the fifth group named ‘short-lived outcrossers with long zygomorphic flowers’ consisted on small perennial and short-lived herbs with long self-compatible zygomorphic flowers that were unable to self-pollinate.

In total, there were 1,126 species of floral visitors with 6,325 interactions recorded with plants. Most plants interacted with bees (2,256 interactions) and non-syrphid Diptera (1,768) followed by syrphids (845), Lepidoptera (437), Coleoptera (432) and non-bee Hymenoptera (362).

### Overall motif patterns

Most motifs were under- or over-represented (close to the 1st and 99th percentile, respectively) in the comparison between empirical and simulated networks (**Figure 2**). Motifs 3, 5, 9, 10 and 14 were underrepresented in empirical networks, that is, all were close to the 1st percentile and under the 25th percentile. Interestingly, four out of five of these motifs belonged to the largest path length classification (i.e., core-peripheral). In addition, motifs 2, 6, 7, 16 and 17 were over-represented, all over the 75th percentile. In contrast to the under-represented motifs, over-represented motifs belonged to the two shortest path length groups (i.e., complete and fan). The remaining motifs (i.e., 4, 8, 11, 12, 13, and 15) were between the 25th and 75th percentile.

**Figure 2.**
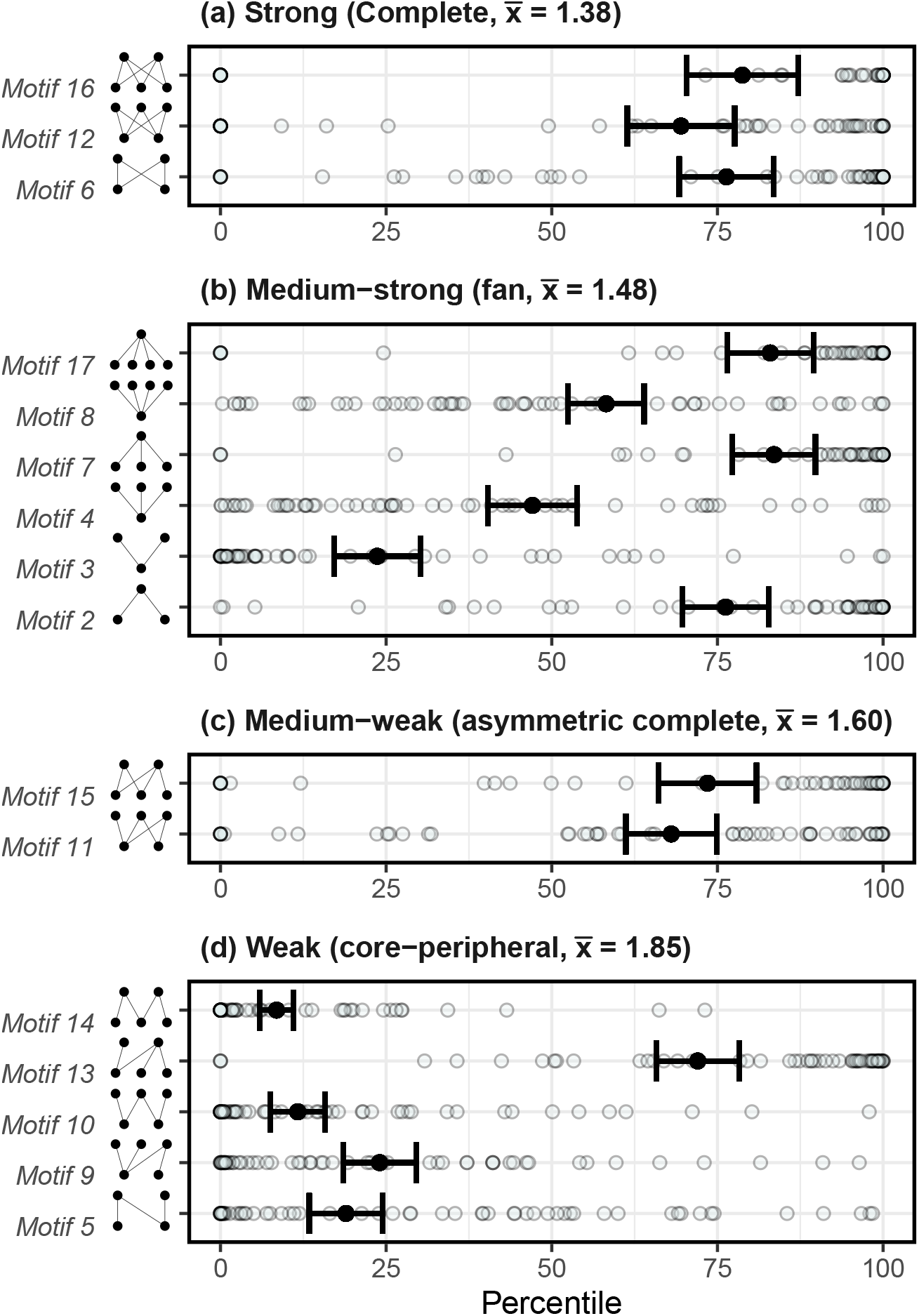
Comparison of motif frequencies between empirical and simulated networks grouped by average path length (plots a, b, c and d) as determined in Simmons et al. (2020). This is shown with the mean percentage of motif frequencies in empirical networks that were over the motif frequencies of the simulated ones (percentiles). This was done by network (light blue transparent dots) and then averaged for all networks (black dots with error bars that correspond to the standard deviation).

### Over- and under-represented groups on motif positions

The comparison of the plant and floral visitor group frequencies per motif position between empirical and simulated networks showed a 6.36% and 25.38% of over- (>75^*th*^) and under-represented (<25th) groups in the different positions, respectively (**Figure 3**). Floral visitors showed a total of 4.55% and 32.58% of over- and under-represented groups in the different positions and plants 8.18% and 18.18% of over- and under-represented groups, respectively. Notably, the differences across groups were more marked for floral visitors than for plant groups (the differences between min and max percentiles per position were generally two-three times larger for floral visitors). From most over- to under-represented floral visitors groups on the different motif positions (indicated with the dendrogram order in **Figure 3**), we found: bees, non-syrphids Diptera, syrphids, Coleoptera, non-bee Hymenoptera and Lepidoptera. Although plant functional groups showed less differences between them, there were also more represented groups than others, thus from most over- to under-represented groups on the different motif positions we found: self incompatible perennials with large flowers, small outcrossing perennials, tall plants with unisexual flowers, selfing herbs and short lived outcrossers with long zygomorphic flowers.

**Figure 3.**
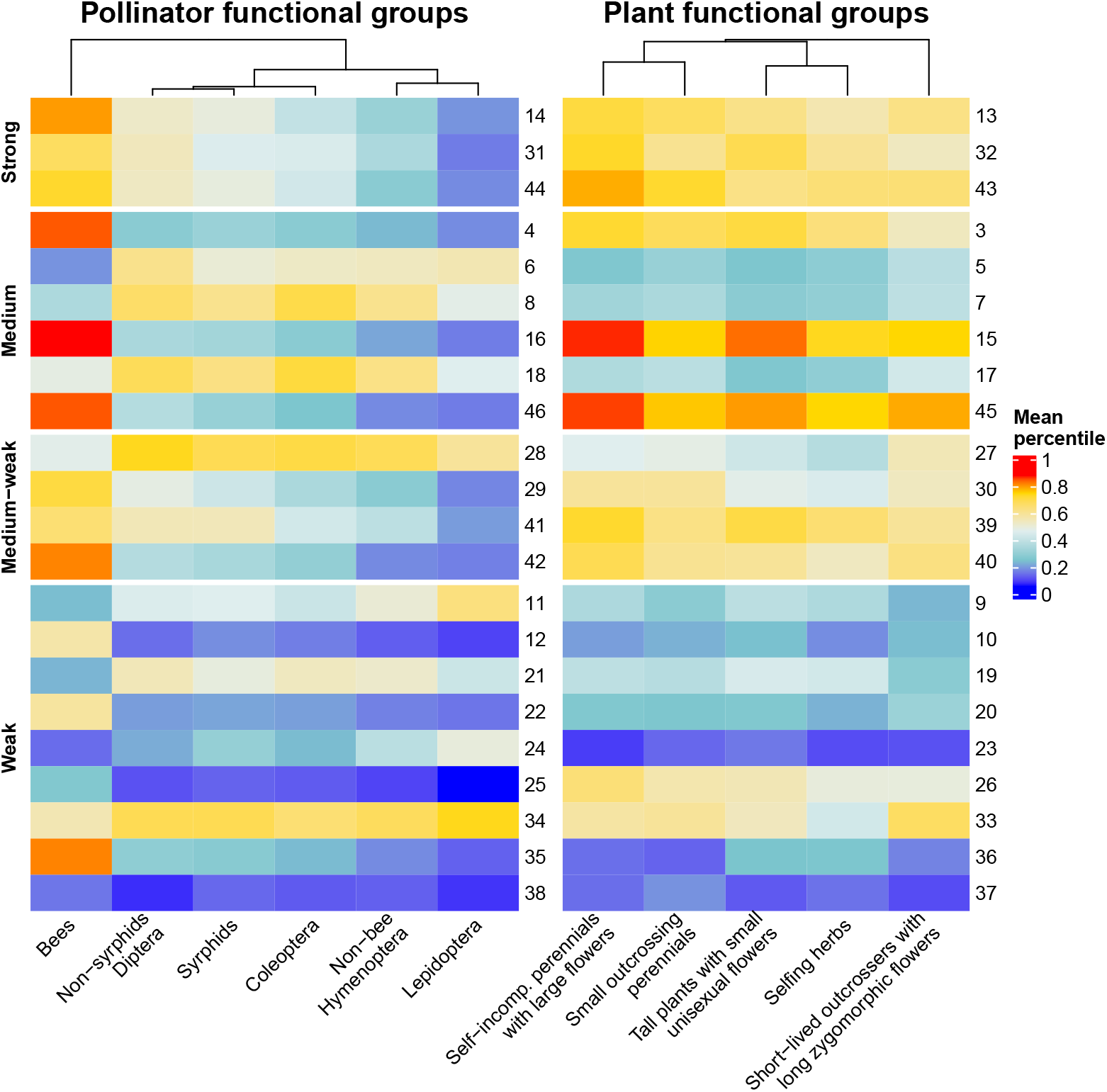
Heatmap indicating under- and over-representation of floral visitor and plant groups in the different motif positions (numbers on the right of each heatmap). The different motif positions are dividied by the average path length clasification determined by Simmons et al. (2020) and they are ordered from the most densely connected motifs (strong or complete) to the less connected ones (weak or core-peripheral). The superior dendrogram indicates the differences across groups with the more separated groups showing larger differences.

For both floral visitor and plant groups we found over- and under-representation on different motif positions that implied different ecological roles. First, for floral visitors, we found that bees were less frequent on motif positions that involved specialization (i.e., lower specificity values) while the rest of taxonomic groups (non-syrphid Diptera, syrphids, Lepidoptera, Coleoptera and non-bee Hymenoptera) were infrequent in generalised motif positions with low number of indirect interactions (**Supporting Information Figure S6 A**). Although plants did not show differences with null models on their frequencies, all plant but tall plants with small unisexual flowers showed a slight tendency to be over-represented on specialised motif positions (**Supporting Information Figure S6 B**). Second, regarding the number of indirect interactions, bees were over-represented in motif positions with low number of indirect interactions (**Supporting Information Figure S7 A**) and the rest of floral visitor groups showed no differences across positions in comparison with simulated networks. For plants, all groups showed a slight trend to be over-represented on motif positions with higher number of indirect interactions (**Supporting Information Figure S7 B**). When we further explore the observed frequencies by number of indirect interactions for floral visitor and plant groups, we found that both bees and tall plants with small unisexual flowers were the only groups with high frequencies on motif positions associated with low number interactions in comparison with the other floral visitor and plant groups (**Supporting Information Figure S8 and Figure S9**).

### Over- and under-represented group combinations of motifs

The different observed frequencies on the different motif positions showed a hierarchical order of probabilities for floral visitors (**Figure 4**). That is, the taxonomic group of ‘bees’ was the most abundant one on all motif positions followed by non-syrphid Diptera which was after bees, the second most frequent group on all positions. Then, the group of syrphids was the most abundant in all positions but two (positions 28 and 31 from motifs 11 and 12, respectively). After syrphids, Coleoptera was the most frequent group on the different positions followed by non-bee Hymenoptera and finally, Lepidoptera which was the less frequent group on all positions but two (positions 12 and 22 from motifs 5 and 9, respectively). Remarkably, plant functional groups were more variable on the different motif positions. However, there were also predominant functional groups found across the different motif positions. The three most frequent groups were ‘tall plants with small unisexual flowers,’ ‘self-incompatible perennials with large flowers’ and ‘small outcrossing perennials.’ On the contrary, the groups of ‘selfing herbs’ and ‘short lived outcrossers with long zygomorphic flowers’ had the lowest probability to be present on the different motifs plant positions.

**Figure 4.**
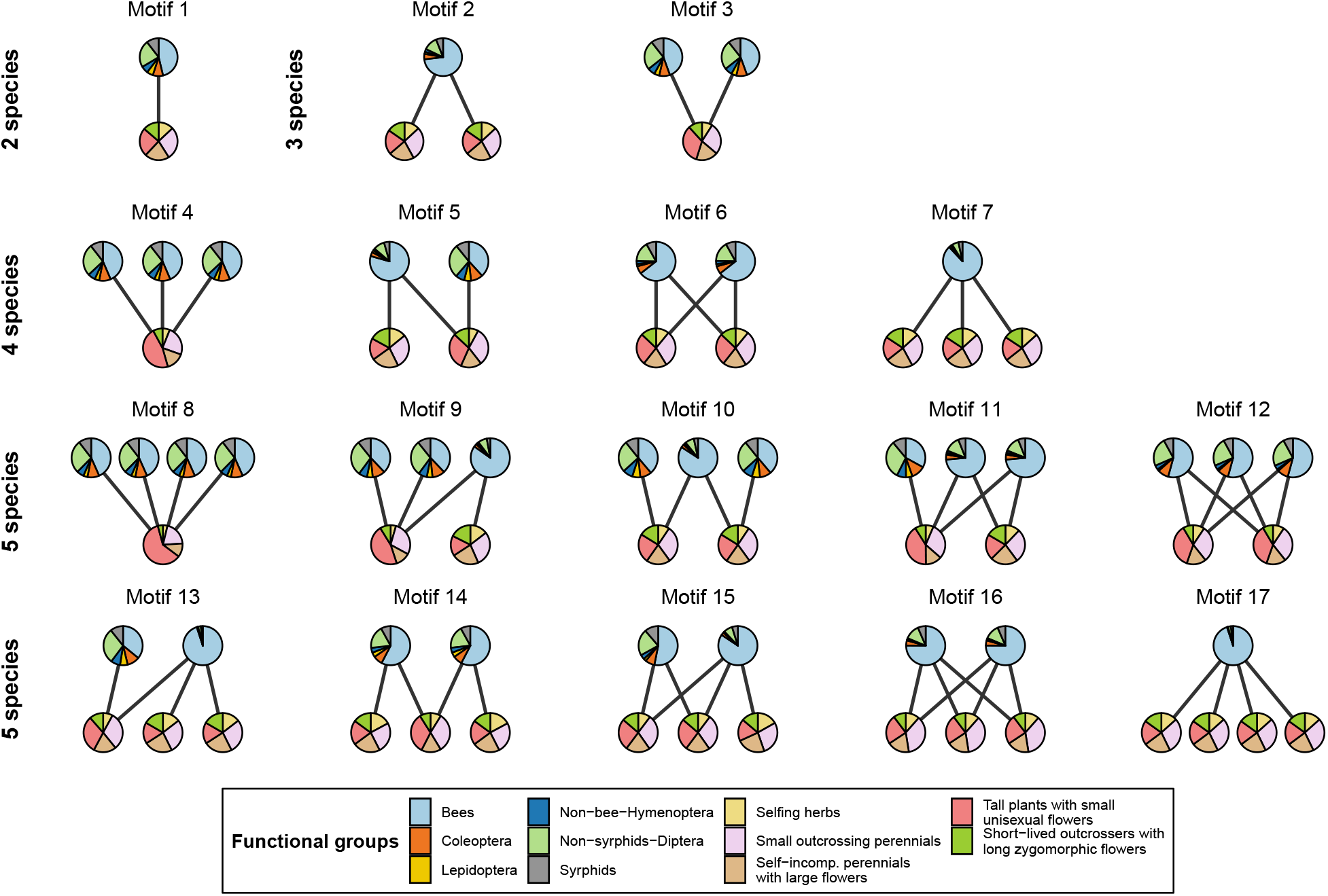
Graphical representation of the probability of finding a given plant and floral visitor group *x* in the position *α* of motif *i, p_i_*(*x, α*), for all the possible motifs from two to five species in bipartite networks. The slices in the nodes for a given functional group *x* are proportional to the corresponding value of *p_i_*(*x, α*).

The statistical comparison with z-scores between the observed and expected plant and floral visitor group combinations showed that we can not recover the observed combinations from its probability of occurrence. In fact, most combinations are under- or over-represented and follow a modified Gaussian distribution (Johnson’s *S_U_*) with 65% of under-represented group combinations, 20% of no statistical difference and 15% of under-represented ones (**Figure 5**). In addition, motifs with small node combinations (i.e., 2 and 3 nodes) only appeared as under-represented. Motifs with 4 and 5 node combinations appeared in the three statistical categories but 4 node motifs had a higher proportion in the under-represented category (10% in comparison with the 1% and the 3% of the no statistical difference and over-represented categories, respectively). Finally, 5 node motifs were in similar proportions in the no statistical difference and over-represented categories (99% and 97%, respectively) but they appeared in lower proportion in the under-represented category (89%). When we explored the identity on the different positions of the most probable motif combinations we found that the most common plant and floral visitor groups were also the most probable on these motif combinations (lower panel **Figure 5**).

**Figure 5.**
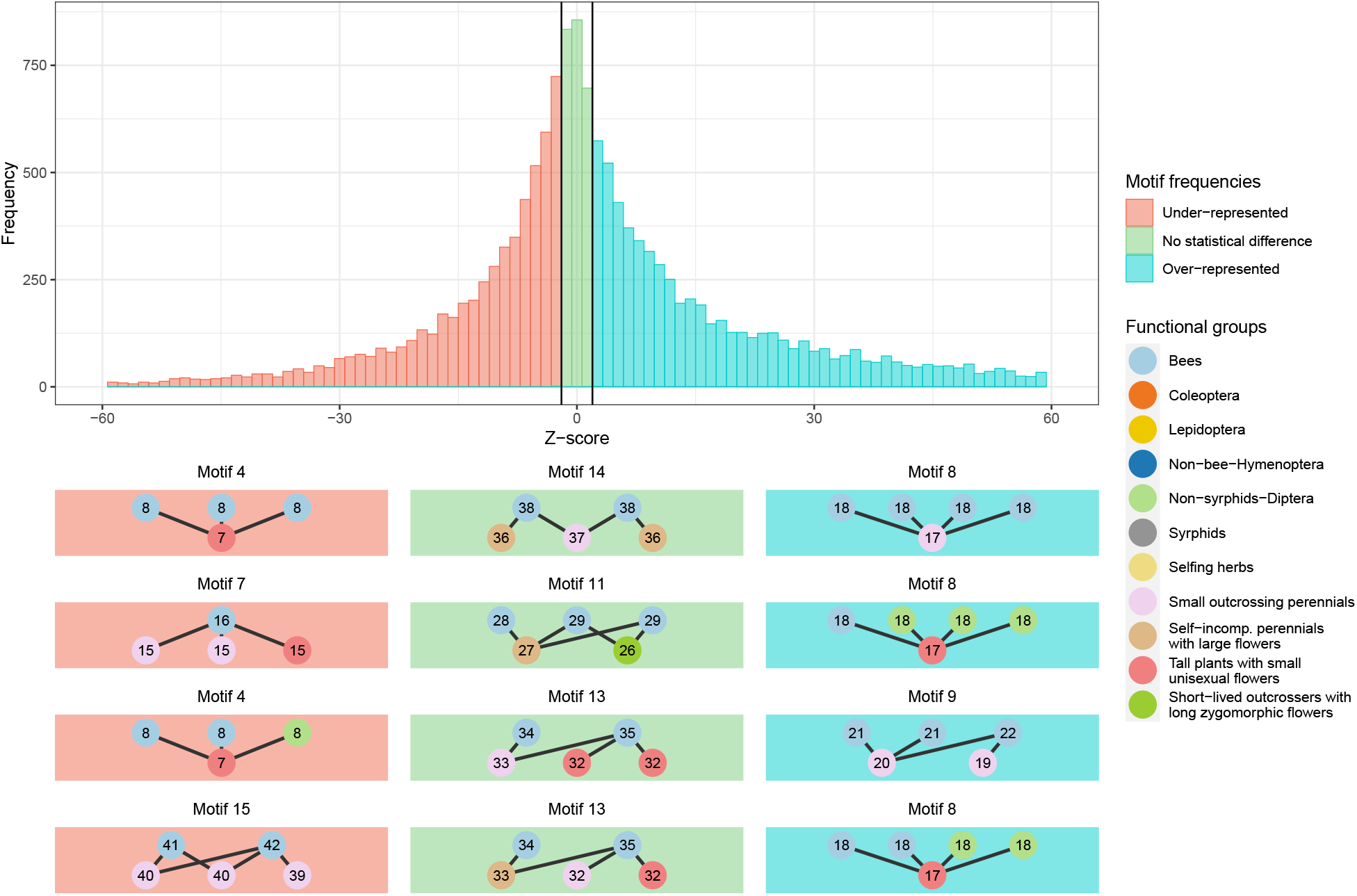
Z-score comparison (95% confidence level) of the observed and expected frequencies of motif combinations with four specific graphical examples of the most probable motifs of being observed for each category delimited by the z-score critical values: under-represented (red), no statistical difference (green) and over-represented (blue).

## Discussion

Understanding the global patterns of network structure is key to unravel the processes that govern community dynamics (Bascompte & Jordano, 2007; Bastolla et al., 2009; Guimarães, 2020). However, common analytical approaches at the node or at the full network level are unable to capture the full complexity of ecological systems (Bergamo et al., 2021; Simmons, Cirtwill, et al., 2019). Because network motifs can capture both direct and indirect interactions, by analysing over- and under-representation of motifs we were able to explore in great detail the structure of interactions of plant-pollinator networks. We have explored for the first time the overall structure and representation of network motifs in real plant-pollinator networks at a macro-ecological scale and shown that the different motifs up to five nodes are consistently over- or under-represented in comparison with simulated networks. Despite the presence of multiple ecological processes (e.g., neutral effects, morphological matching or phenological overlap) shaping motif structure, we found a tendency for over-represented structures with more densely connected motifs (i.e., complete, fan and asymmetric complete). As suggested in Simmons et al. (2020), a higher proportion of these densely connected motifs could reflect that neutral effects are the predominant ecological process governing the structure of network motifs. Thus, as indicated for other network properties like nestedness at the macro-scale level (Krishna et al., 2008; Suweis et al., 2013), or interaction probability at the species level (Bartomeus et al., 2016), species relative abundances (i.e., neutral effects) may be also the main driver governing the meso-scale structure on plant-pollinator networks.

Overall, generalist network motifs for floral visitors were consistently over-represented (e.g., fan-type motifs; **Figure 2**). For instance, when we compare motifs with equal structure for both trophic levels, we found that floral visitors had always over-representation in the positions that implied the highest number of direct links (e.g., positions 16 and 46 from motifs 7 and 17, respectively). Despite this being in good agreement with the current view of the generalist nature of pollinators (Olesen & Jordano, 2002; Ollerton, 2017; Waser et al., 1996), we found that this over-representation of generalist network motifs was driven by the taxonomic group of bees while the rest of the taxonomic groups tended to be under-represented (**Supporting Information Figure S6 A**). In contrast, non-bee groups were over-represented in motif positions associated with specialised roles. This large effect of bees on the overall patterns is no surprise, as bees were the group with the largest number of interaction records in the whole set of networks as also shown for other broad scale plant-pollinator network studies (Carvalheiro et al., 2014; Ollerton, 2017). Interestingly, bees also were the group with lower number of indirect interactions while the rest of floral visitor groups showed clearly higher number of indirect interactions indicating that these non-bee floral visitor groups may experience higher competitive interactions for floral resources (Thomson & Page, 2020). Despite plant functional groups showed little differences between them on their motif frequencies, all groups showed a tendency to be over-represented on motif positions that were associated with higher levels of specialisation. This tendency towards specialist roles may be the result of co-evolutionary processes (e.g., trait-matching) between plants and pollinators to optimise pollen transfer (Moreira-Hernández & Muchhala, 2019). Remarkably, only tall plants with small unisexual flowers showed higher frequencies on motifs that were associated with low number of indirect interactions with likely direct implications on pollen transfer for this group that needs to guarantee cross-pollination.

The probabilities of appearance of each functional group on the different motif positions were insufficient to predict most of the observed network position combinations (**Figure 5**). This implies that not just the representation of network motifs types but also the identity of the elements of their structure is non-random. Notably, we found that just 30% of the possible 53,250 motif combinations are realized. Thus, similarly to the presence of forbidden links at the network level (Jordano, 2016; Olesen et al., 2011), we also found the existence of forbidden motif positions and those represented almost two thirds of the total possible motif combinations. This concurs well with the general idea that plant-pollinator networks have a high proportion of non-realized direct interactions (Bascompte & Jordano, 2013; Jordano et al., 2003), and as a result, also a high proportion of non-realized combinations of indirect interactions. Some of this non-observed direct interactions are probably due to under-sampling (Dorado et al., 2011; Jordano, 2016). However, a large fraction of the possible interactions do not take place because of life-history constraints such as species co-occurrence, trait matching processes and phenological uncoupling (Bartomeus et al., 2016; Jordano, 2016; Olesen et al., 2011). Despite the advances in the understanding on the ecological processes that influence species pairwise interactions (Bartomeus et al., 2016; Peralta et al., 2020), we still know little about how these processes determine competitive and facilitative interactions in mutualistic networks. Thus, as suggested by Simmons et al. (2020), exploring the proportion of these realized network motifs that are competitive or facilitative is a necessary next step to improve our understanding of the architecture of biodiversity (Bascompte & Jordano, 2007).

Despite the obvious usefulness of functional groups in summarizing species ecological roles (e.g., similar reproductive biology for plants and behaviour for pollinators), delimiting species into functional groups is often not trivial. For instance, taxonomic diversity and trait availability limit our ability to properly represent the wide range of reproductive strategies (Dellinger, 2020) and different clustering analysis can lead to distinct outcomes (e.g., k-means and hierarchical clustering). Further, here we divided floral visitors on the taxonomic rank level because it represented adequately pollinator ecological roles without the tremendous effort of compiling trait data for over 1000 species of floral visitors. However, further groupings with traits could help revealing greater relevance of trait-matching processes on indirect interactions. We believe that this could be investigated by exploring which proportion of over- and under-represented motifs present high and low trait-matching, respectively. Moreover, we analysed network motifs focusing exclusively on presence-absence interactions which assume that all interactions are equally important as suggested by Simmons et al. (2020), further work should investigate how to account for the quantitative measurements of the strength of plantpollinator interactions. Finally, despite the wide geographical coverage of our plant-pollinator dataset, there are major gaps for some regions of the world with a clear bias towards Europe, America and islands which have long interested ecologist (MacArthur & Wilson, 1963; Warren et al., 2015). These biases are also found in global datasets of networks where the data is highly fragmented across space (Poisot et al., 2021) and more efforts are needed to capture global patterns of plant-pollinator interactions.

In conclusion, by following the framework of Simmons et al. (2020) we have been able to explore competitive and facilitative interactions in plant-pollinator networks worldwide and found that indirect interactions are non-random and are more strongly related to neutral processes than to any other ecological process. Further, we found that plant and floral visitor groups are associated with different motif positions that involve distinct ecological roles (e.g., generalisation versus specialisation) and that the different motif combinations are strongly constrained. Although we are still far to predict how species interact, our work is a first step to elucidate beyond pairwise interactions.

## Acknowledgements

This study was supported by the European project SAFEGUARD (101003476 H2020-SFS-2019-2). JBL also thanks the University of New England for the funding provided to carry out this work. Finally, we would like to thank David García-Callejas for his useful comments on the manuscript before submission.

## Author contributions

IB, AA-P and JBL designed the idea. JBL collected the data. AA-P lead the analysis with contributions of JBL and guidance of IB. JBL wrote the manuscript with contributions of IB and AA-P.

