## Supporting Information for "The non-random assembly of functional motifs in plant-pollinator networks"

**TABLES**

**Table S1** List of plant-pollinator network studies.

**Table S2** List of the plant species used in this study.

**Table S3** Type of plant traits used in this study.

**FIGURES**

**Figure S1** Map with the plant-pollinator study locations.

**Figure S2** Over and under-estimated motif frequencies.

**Figure S3** Over and under-estimated groups on motif positions.

**Figure S4** Plant trait composition of functional groups.

**Figure S5** Dendrogram with plant functional groups.

**Figure S6** Association between the percentile and specificity of the different plant and floral visitor groups on motif positions.

**Figure S7** Association between the percentile and number of indirect interactions of the different plant and floral visitor groups on motif positions.

**Figure S8** Frequency of floral visitor groups and number of indirect interactions per motif positions.

**Figure S9** Frequency of plant groups and number of indirect interactions per motif positions.

**Table S1.** List of studies ordered by author with the year of publication, number of contributed networks and digital object identifier.

| First author | Year | Number of networks | Country | DOI |
|---|---|---|---|---|
| Arroyo-Correa | 2019 | 3 | New Zealand | https://doi.org/10.1111/1365-2745.13332 |
| Bartomeus | 2008 | 6 | Spain | https://doi.org/10.1007/s00442-007-0946-1 |
| Bartomeus | 2015 | 16 | Spain | https://github.com/ibartomeus/BeeFunData |
| Bundgaard | 2003 | 1 | Denmark | Unpublished, Master thesis |
| Burkle | 2013 | 1 | United States | https://doi.org/10.1126/science.1232728 |
| Dicks | 2002 | 2 | England | https://doi.org/10.1046/j.0021-8790.2001.00572.x |
| Dupont | 2003 | 3 | Denmark | https://doi.org/10.1111/j.1365-2656.2008.01501.x |
| Elberling | 1999 | 1 | Sweden | https://doi.org/10.1111/j.1600-0587.1999.tb00507.x |
| Fang | 2008 | 1 | China | https://doi.org/10.1111/1749-4877.12190 |
| Inouye | 1988 | 1 | United States | https://doi.org/10.1111/j.1442-9993.1988.tb00968.x |
| Kaiser-Bunbury | 2017 | 8 | Seychelles | https://doi.org/10.1038/nature21071 |
| Kaiser-Bunbury | 2011 | 6 | Seychelles | https://doi.org/10.1111/j.1365-2745.2010.01732.x |
| Kaiser-Bunbury | 2010 | 2 | Mauritius | https://doi.org/10.1016/j.ppees.2009.04.001 |
| Lundgren | 2005 | 1 | Denmark (Greenland) | https://doi.org/10.1657/1523-0430(2005)037[0514:TDAHCW]2.0.CO;2 |
| Olesen | 2002 | 2 | Mauritius and Portugal (Azores) | https://doi.org/10.1046/j.1472-4642.2002.00148.x |
| Peralta | 2006 | 4 | Argentina | https://doi.org/10.1111/ele.13510 |
| Small | 1976 | 1 | Japan | /13960/t4km08d21 |
| Souza | 2017 | 1 | Brazil | https://doi.org/10.1111/1365-2745.12978 |

**Table S2.** List of the 503 plant species used in this study.

| List of plant species | | |
|---|---|---|
| *Acantholippia seriphioides* | *Erica umbellata* | *Peponidium carinatum* |
| *Achillea millefolium* | *Erigenia bulbosa* | *Pereskia sacharosa* |
| *Achillea wilsoniana* | *Erophaca baetica* | *Persicaria runcinata* |
| *Aciphylla glacialis* | *Eryngium campestre* | *Persicaria vivipara* |
| *Aciphylla simplicifolia* | *Erythronium albidum* | *Phlomis atropurpurea* |
| *Acrothamnus montanus* | *Erythrospermum monticolum* | *Phlomis purpurea* |
| *Actaea yunnanensis* | *Erythroxylum macrocarpum* | *Phlomis tatsienensis* |
| *Adenophora capillaris* | *Erythroxylum sechellarum* | *Phlox divaricata* |
| *Adenophora jasionifolia* | *Eugenia kanakana* | *Phoenicophorium borsigianum* |
| *Adenophora khasiana* | *Eugenia orbiculata* | *Phyllanthus phillyreifolius* |
| *Aeschynomene mollicula* | *Euphorbia pyrifolia* | *Picris hieracioides* |
| *Agarista salicifolia* | *Euphorbia segetalis* | *Pilosella officinarum* |
| *Agrimonia pilosa* | *Euphrasia collina* | *Pimelea ligustrina* |
| *Ajuga forrestii* | *Euphrasia regelii* | *Pinguicula alpina* |
| *Allium ampeloprasum* | *Excoecaria benthamiana* | *Pittosporum senacia* |
| *Allium atrosanguineum* | *Faujasiopsis flexuosa* | *Plantago lanceolata* |
| *Allium cyathophorum* | *Filipendula ulmaria* | *Pleurospermum davidii* |
| *Allium wallichii* | *Flagellaria indica* | *Pleurostylia leucocarpa* |
| *Aloysia gratissima* | *Freesia refracta* | *Polemonium reptans* |
| *Alstonia macrophylla* | *Gaertnera petrinensis* | *Polyscias mauritiana* |
| *Anacamptis morio* | *Gaertnera psychotrioides* | *Portulaca fluvialis* |
| *Anaphalioides alpinum* | *Gaertnera rotundifolia* | *Potentilla crantzii* |
| *Anaphalis nepalensis* | *Galactites tomentosus* | *Potentilla erecta* |
| *Anchusa azurea* | *Galeopsis bifida* | *Potentilla fallens* |
| *Andromeda glaucophylla* | *Galium saxatile* | *Potentilla griffithii* |
| *Andryala integrifolia* | *Galium verum* | *Potentilla lancinata* |
| *Anemone rivularis* | *Gastonia crassa* | *Potentilla rubricaulis* |
| *Anemonella thalictroides* | *Gaylussacia baccata* | *Premna serratifolia* |
| *Angelica archangelica* | *Geniostoma borbonicum* | *Primula poissonii* |
| *Anthriscus sylvestris* | *Genista anglica* | *Primula secundiflora* |
| *Antidesma madagascariense* | *Genista hirsuta* | *Primula veris* |
| *Antirhea borbonica* | *Genista pilosa* | *Prosopis flexuosa* |
| *Aphloia theiformis* | *Gentiana chungtienensis* | *Prosopis rubriflora* |
| *Arabis alpina* | *Gentiana crassicaulis* | *Prostanthera cuneata* |
| *Arachis microsperma* | *Gentianella diemensis* | *Prunella vulgaris* |
| *Arctium lappa* | *Geranium delavayi* | *Pseudognaphalium affine* |
| *Arctotheca calendula* | *Geranium maculatum* | *Psiadia terebinthina* |
| *Arenaria yunnanensis* | *Glionnetia sericea* | *Psidium cattleyanum* |
| *Argusia argentea* | *Gomphrena celosioides* | *Psychotria pervillei* |
| *Armeria velutina* | *Gomphrena elegans* | *Psychotria terniflora* |
| *Arnica montana* | *Grangeria borbonica* | *Pterocephalus hookeri* |
| *Aronia melanocarpa* | *Halenia elliptica* | *Pyrola grandiflora* |
| *Asphodelus fistulosus* | *Halimium commutatum* | *Pyrostria bibracteata* |
| *Aster diplostephioides* | *Halimium halimifolium* | *Pyrostria fasciculata* |
| *Aster oreophilus* | *Hancea integrifolia* | *Ranunculus acris* |
| *Aster souliei* | *Harrimanella hypnoides* | *Ranunculus hispidus* |
| *Aster vestitus* | *Harungana madagascariensis* | *Reseda luteola* |

*Aster yunnanensis*
*Astragalus alpinus*
*Astragalus pullus*
*Atamisquea emarginata*
*Ayenia tomentosa*
*Azorina vidalii*
*Badula insularis*
*Badula platyphylla*
*Baeckea gunniana*
*Bakerella hoyifolia*
*Bartsia alpina*
*Bertiera zaluzania*
*Beta vulgaris*
*Bistorta macrophylla*
*Bistorta sinomontana*
*Bituminaria bituminosa*
*Brassica fruticulosa*
*Buddleja mendozensis*
*Calendula arvensis*
*Calluna vulgaris*
*Calophyllum eputamen*
*Calopogon tuberosus*
*Camassia scilloides*
*Campanula giesekiana*
*Campanula rotundifolia*
*Campnosperma seychellarum*
*Camptosema paraguariense*
*Canthium bibracteatum*
*Cardamine concatenata*
*Carduus crispus*
*Carpobrotus acinaciformis*
*Casearia coriacea*
*Cassiope tetragona*
*Castela coccinea*
*Celmisia asteliifolia*
*Celmisia graminifolia*
*Centaurea nigra*
*Cerastium alpinum*
*Cerastium fontanum*
*Cereus hildmannianus*
*Chamaedaphne calyculata*
*Chamomilla suaveolens*
*Chassalia coriacea*
*Chassalia petrinensis*
*Chrysobalanus icaco*
*Cinnamomum verum*
*Cirsium arvense*
*Cirsium eriophoroides*
*Cirsium palustre*
*Cirsium pratense*

*Helenium donianum*
*Helichrysum proteoides*
*Helichrysum stoechas*
*Herissantia nemoralis*
*Herpetospermum pedunculosum*
*Heteropterys glabra*
*Hibiscus tiliaceus*
*Hieracium praealtum*
*Hieracium umbellatum*
*Homalanthus populifolius*
*Hydrophyllum appendiculatum*
*Hydrophyllum virginianum*
*Hypericum perfolatum*
*Hypericum perforatum*
*Hypochaeris radicata*
*Impatiens chungtienensis*
*Ipomoea macrantha*
*Isostigma hoffmannii*
*Ixeridium biparum*
*Ixora parviflora*
*Ixora pudica*
*Juncus allioides*
*Kalmia angustifolia*
*Kalmia polifolia*
*Kunzea muelleri*
*Labourdonnaisia calophylloides*
*Larrea divaricata*
*Larrea nitida*
*Lathyrus clymenum*
*Lathyrus pratensis*
*Lavandula pedunculata*
*Lavandula stoechas*
*Lecanophora heterophylla*
*Ledum palustre*
*Leontodon saxatilis*
*Lepidaploa pseudomuricata*
*Leptospermum scoparium*
*Leucaena leucocephala*
*Leucanthemum vulgare*
*Leucochrysum albicans*
*Ligularia cymbulifera*
*Ligularia dictyoneura*
*Ligularia lankongensis*
*Lilium duchartrei*
*Linaria viscosa*
*Linum bienne*
*Lippia alba*
*Lobularia maritima*
*Lotus corniculatus*
*Lotus pedunculatus*

*Rhododendron lapponicum*
*Richea continentis*
*Rosa rubiginosa*
*Roscheria melanochaetes*
*Rosmarinus officinalis*
*Roussea simplex*
*Rubus alceifolius*
*Rubus plicatus*
*Ruellia tweediana*
*Ruta chalepensis*
*Salix fragilis*
*Salix lanata*
*Salix polaris*
*Salix repens*
*Salix reticulata*
*Salvia przewalskii*
*Sanicula odorata*
*Saussurea wardii*
*Saxifraga aizoides*
*Saxifraga oppositifolia*
*Scabiosa atropurpurea*
*Scabiosa stellata*
*Scaevola taccada*
*Schinus fasciculata*
*Scoparia montevidensis*
*Scorzonera humilis*
*Sedum roseum*
*Sedum sediforme*
*Senecio lautus*
*Senecio pinnatus*
*Senna aphylla*
*Sida rhombifolia*
*Sideroxylon cinereum*
*Sideroxylon puberulum*
*Silene acaulis*
*Silene dioica*
*Silene flos-cuculi*
*Silene gracilicaulis*
*Silene suecica*
*Silene vulgaris*
*Silene yunnanensis*
*Smilax anceps*
*Solidago sempervirens*
*Solidago virgaurea*
*Sonchus tenerrimus*
*Soulamea terminaloides*
*Spartium junceum*
*Spenceria ramalana*
*Spermacoce eryngioides*
*Spiraea alba*

*Cirsium vulgare*
*Cistus albidus*
*Cistus crispus*
*Cistus ladanifer*
*Cistus libanotis*
*Cistus monspeliensis*
*Cistus salviifolius*
*Claoxylon linostachys*
*Claytonia virginica*
*Cleistocactus baumannii*
*Clematis akebioides*
*Clematis campestris*
*Cleome guianensis*
*Clinopodium repens*
*Cnidoscolus urens*
*Coccoloba guaranitica*
*Codonopsis convolvulacea*
*Coffea macrocarpa*
*Coffea mauritiana*
*Colea seychellarum*
*Condalia microphylla*
*Convolvulus althaeoides*
*Convolvulus arvensis*
*Corokia cotoneaster*
*Craterispermum microdon*
*Crepis capillaris*
*Crithmum maritimum*
*Croton fothergillifolius*
*Croton grangerioides*
*Cuphea thymoides*
*Cyananthus delavayi*
*Cynoglossum amabile*
*Daucus carota*
*Deckenia nobilis*
*Delphinium tricorne*
*Dianthus armeria*
*Dianthus caryophyllus*
*Diapensia lapponica*
*Dicentra cucullaria*
*Digitalis purpurea*
*Dillenia ferruginea*
*Dillenia suffruticosa*
*Diospyros revaughanii*
*Diospyros seychellarum*
*Diplotaxis virgata*
*Dipsacus asperoides*
*Dipsacus chinensis*
*Dodonaea viscosa*
*Doratoxylon apetalum*
*Dorycnium pentaphyllum*

*Lupinus angustifolius*
*Lycium chilense*
*Lysimachea europaea*
*Maianthemum racemosum*
*Maianthemum trifolium*
*Malva multiflora*
*Medusagyne oppositifolia*
*Melampyrum pratense*
*Melicope chapelieri*
*Melicope lunu-ankenda*
*Memecylon eleagni*
*Memecylon ovatifolium*
*Menodora decemfida*
*Mentha canadensis*
*Mertensia virginica*
*Microseris lanceolata*
*Microtea scabrida*
*Microula sikkimensis*
*Mimosa hexandra*
*Mimosa sensibilis*
*Mimulus moschatus*
*Mimusops erythroxylon*
*Mimusops sechellarum*
*Molinaea arborea*
*Molinaea macrantha*
*Mollugo verticillata*
*Monarda bradburiana*
*Morinda citrifolia*
*Muehlenbeckia axillaris*
*Muehlenbeckia complexa*
*Murdannia nudiflora*
*Mycelis muralis*
*Myosotis alpestris*
*Nematolepis ovatifolia*
*Nemopanthus mucronatus*
*Nepenthes pervillei*
*Nepeta stewartiana*
*Nephrosperma vanhoutteanum*
*Neptunia plena*
*Northia seychellana*
*Ochna kirkii*
*Ochna mauritiana*
*Ocotea laevigata*
*Olea lancea*
*Olearia bullata*
*Onosma confertum*
*Opuntia elatior*
*Opuntia stricta*
*Opuntia sulphurea*
*Origanum vulgare*

*Stachys sylvatica*
*Stachytarpheta indica*
*Stachytarpheta jamaicensis*
*Stellaria graminea*
*Stellaria yunnanensis*
*Stillingia lineata*
*Stylosanthes hamata*
*Succisa pratensis*
*Suriana maritima*
*Swertia forrestii*
*Syzygium commersonii*
*Syzygium coriaceum*
*Syzygium glomeratum*
*Syzygium jambos*
*Syzygium mauritianum*
*Syzygium petrinense*
*Syzygium venosum*
*Syzygium wrightii*
*Tabernaemontana persicarifolia*
*Talinum fruticosum*
*Tambourissa peltata*
*Taraxacum campylodes*
*Teucrium fruticans*
*Thalictrum delavayi*
*Thalictrum rostellatum*
*Thapsia villosa*
*Thespesia populnea*
*Thymelaea hirsuta*
*Thymus mastichina*
*Tibetia himalaica*
*Tillandsia didisticha*
*Timonius sechellensis*
*Toddalia asiatica*
*Torilis japonica*
*Tradescantia virginiana*
*Trifolium arvense*
*Trifolium dubium*
*Trifolium pratense*
*Trifolium repens*
*Tripogandra glandulosa*
*Tristemma mauritianum*
*Trochetia blackburniana*
*Trollius europaeus*
*Trollius vaginatus*
*Tuberaria guttata*
*Turnera angustifolia*
*Turraea rigida*
*Ulex parviflorus*
*Urospermum picroides*
*Uvularia grandiflora*

| | | |
|---|---|---|
| *Dracaena concinna* | *Orites lancifolius* | *Vaccinium myrtilloides* |
| *Dracaena reflexa* | *Oxalis violacea* | *Vaccinium myrtillus* |
| *Dryas integrifolia* | *Oxylobium ellipticum* | *Vaccinium uliginosum* |
| *Dryas octopetala* | *Ozothamnus leptophyllus* | *Vaccinium vitis-idaea* |
| *Dubyaea bhotanica* | *Pandanus barkleyi* | *Verbascum thapsus* |
| *Echinopsis candicans* | *Pandanus rigidifolius* | *Veronica brachysiphon* |
| *Echinopsis rhodotricha* | *Pandanus wiehei* | *Vicia cracca* |
| *Echium plantagineum* | *Paragenipa wrightii* | *Vicia lutea* |
| *Echium sabulicola* | *Paraserianthes serratifolia* | *Vicia sativa* |
| *Eleutherine bulbosa* | *Parkinsonia praecox* | *Viola biflora* |
| *Embelia angustifolia* | *Parnassia palustris* | *Viola pubescens* |
| *Empetrum nigrum* | *Passiflora suberosa* | *Viola sororia* |
| *Enemion biternatum* | *Pavonia sidifolia* | *Wahlenbergia albomarginata* |
| *Epacris petrophila* | *Pedicularis cephalantha* | *Wahlenbergia ceracea* |
| *Epilobium angustifolium* | *Pedicularis densispica* | *Waltheria indica* |
| *Epilobium gunnianum* | *Pedicularis dichotoma* | *Warneckea trinervis* |
| *Epilobium hirsutum* | *Pedicularis rex* | *Wikstroemia indica* |
| *Epilobium latifolium* | *Pedicularis siphonantha* | *Xylopia lamarckii* |
| *Epilobium tibetanum* | *Pedicularis tricolor* | *Ziziphus mistol* |
| *Erica scoparia* | *Pemphis acidula* | *Zuccagnia punctata* |
| *Erica tetralix* | *Pentachondra pumila* | |

**Table S3.** Traits used to delimit the different plant functional groups divided in quantitative and categorical traits.

| Quantitative traits | | Categorical traits | |
|---|---|---|---|
| **Type** | **Traits** | **Type** | **Traits** |
| **Vegetative** | Plant height (m) | Vegetative | **Lifepan** |
| **Floral** | Flower width (mm) | Vegetative | **Life form** |
| **Floral** | Flower length (mm) | Floral | **Flower shape** |
| **Floral** | Inflorescence width (mm) | Floral | **Flower symmetry** |
| **Floral** | Style length (mm) | Reproductive | **Autonomous selfing** |
| **Floral** | Ovules per flower | Reproductive | **Compatibility system** |
| **Floral** | Flowers per plant | Reproductive | **Breeding system** |
| **Reproductive** | Autonomous selfing (fruit set) | | |

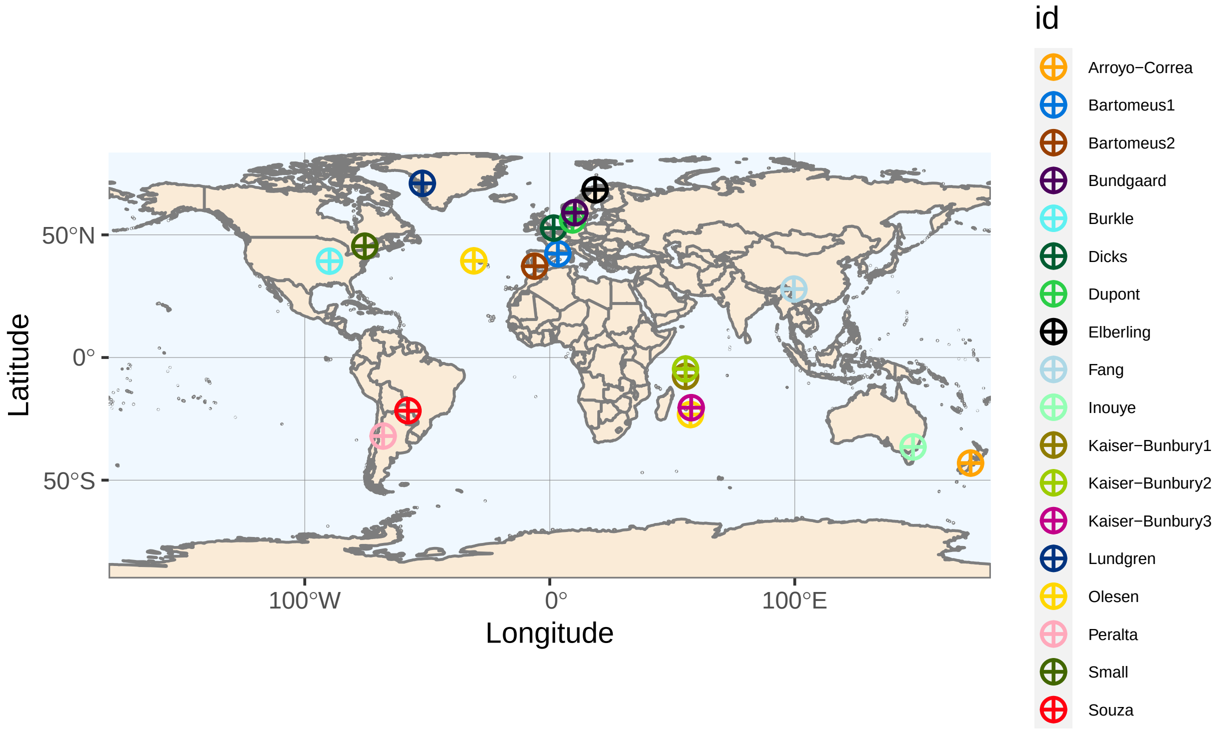


**Figure S1.** Map with the different locations of the plant-pollinator studies used in this work to explore woldwide patterns at the meso-scale level. For each study just one location is shown except for the study of Olesen that was conducted in two very different locations (Mauritius and Azores).

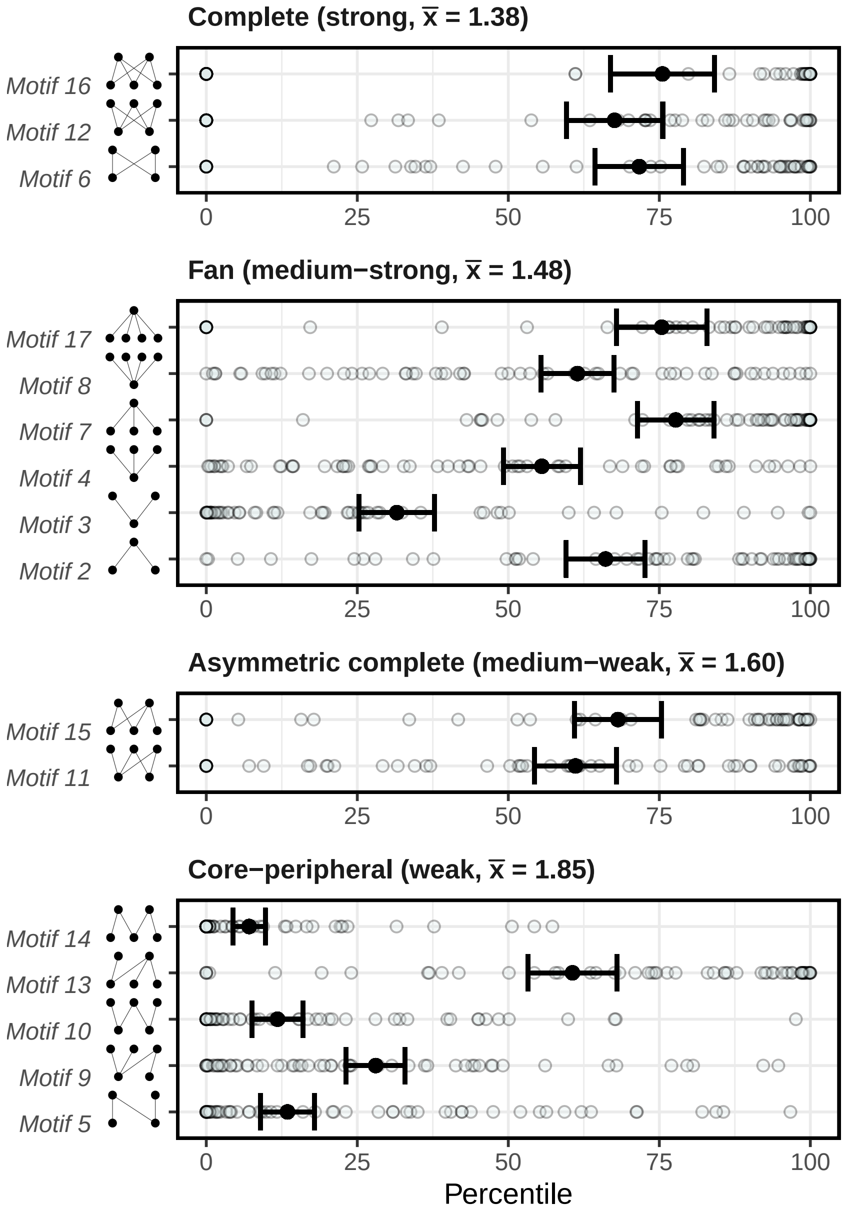


**Figure S2.** Comparison of motif frequencies between empirical and simulated networks grouped by average path length (plots a, b, c and d) as determined in Simmons et al. (2020) without considering singletones. This is shown with the mean percentage of motif frequencies in empirical networks that were over the motif frequencies of the simulated ones (percentiles). This was done by network (light blue dots) and then averaged for all networks (black dots with error bars that correspond to the standard deviation).

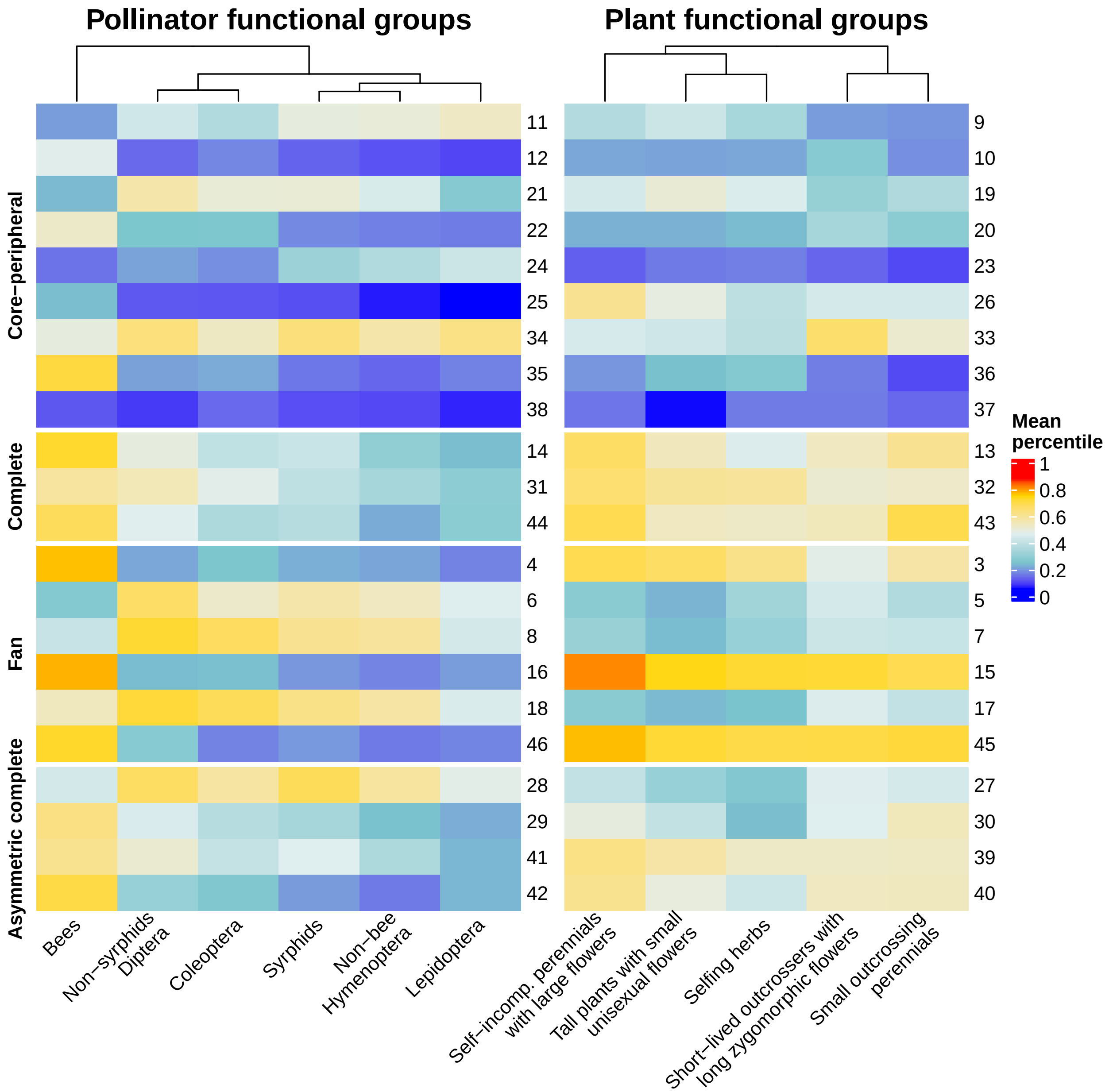


**Figure S3.** Heatmap indicating under- and over- representation of pollinator and plant functional groups in the different motif positions after removing non-robust links (singletones). The different motif positions are dividied by the average path length clasification determined by Simmons et al. (2020). The superior dendrogram indicates the differences across groups with the more separated groups showing larger differences.

### Plant functional group composition

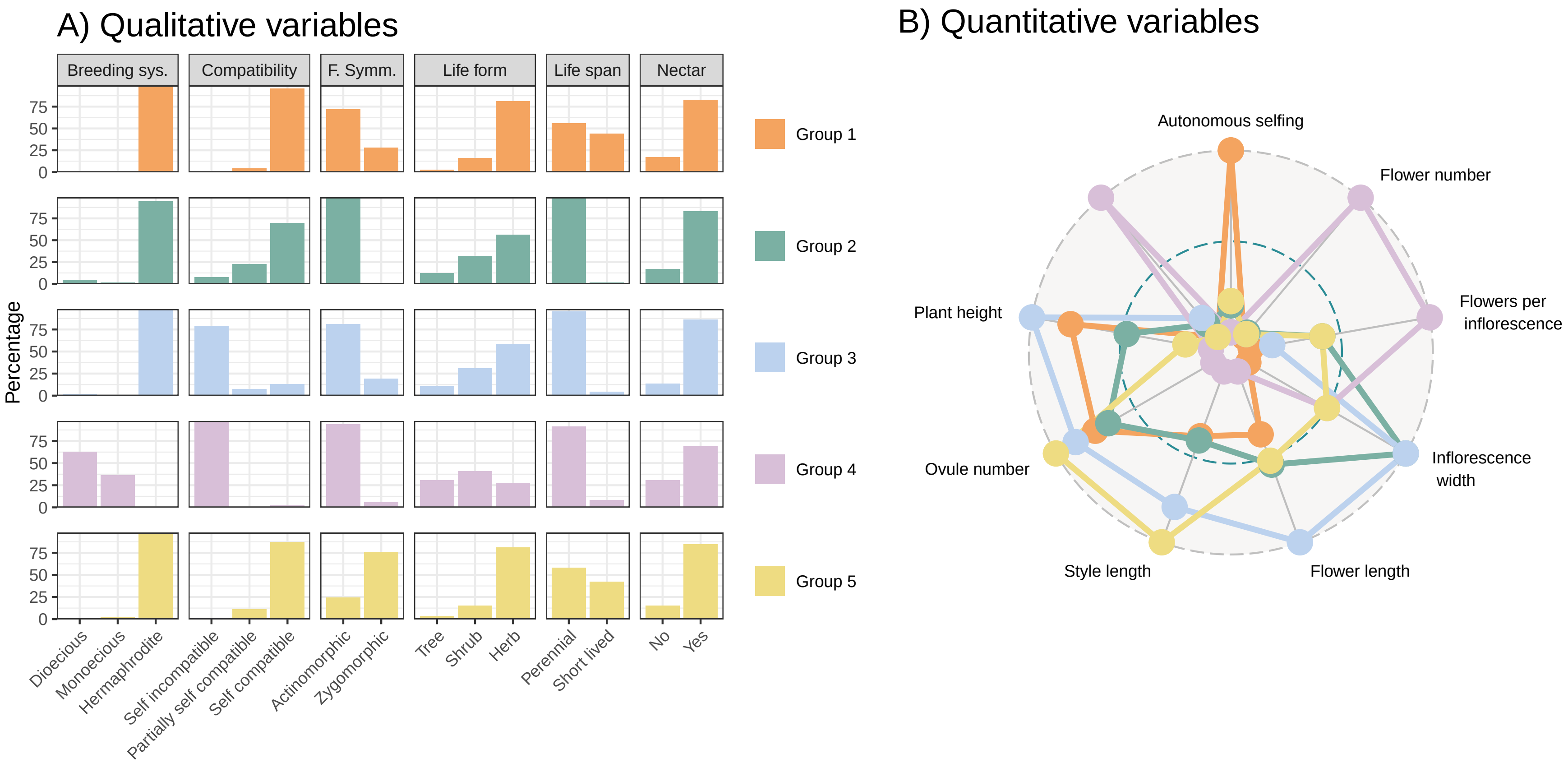


**Figure S4.** Plant functional group composition separated in qualitative and quantitative variables. Panel A) shows the percentage of the different categories within trait represented with different colours for each functional group. Plot B) shows the radar plot of the different quantitative variables standardise on the same scale also coloured with the same patterns of colours as qualitative variables per cluster or functional group. Group 1 corresponds to short-lived selfers; group 2 to small outcrossing perennials; group 3 to self-incompatible perennials with large flowers; group 4 to tall plants with small unisexual flowers; and, group 5 to short-lived outcrossers with long zygomorphic flowers.

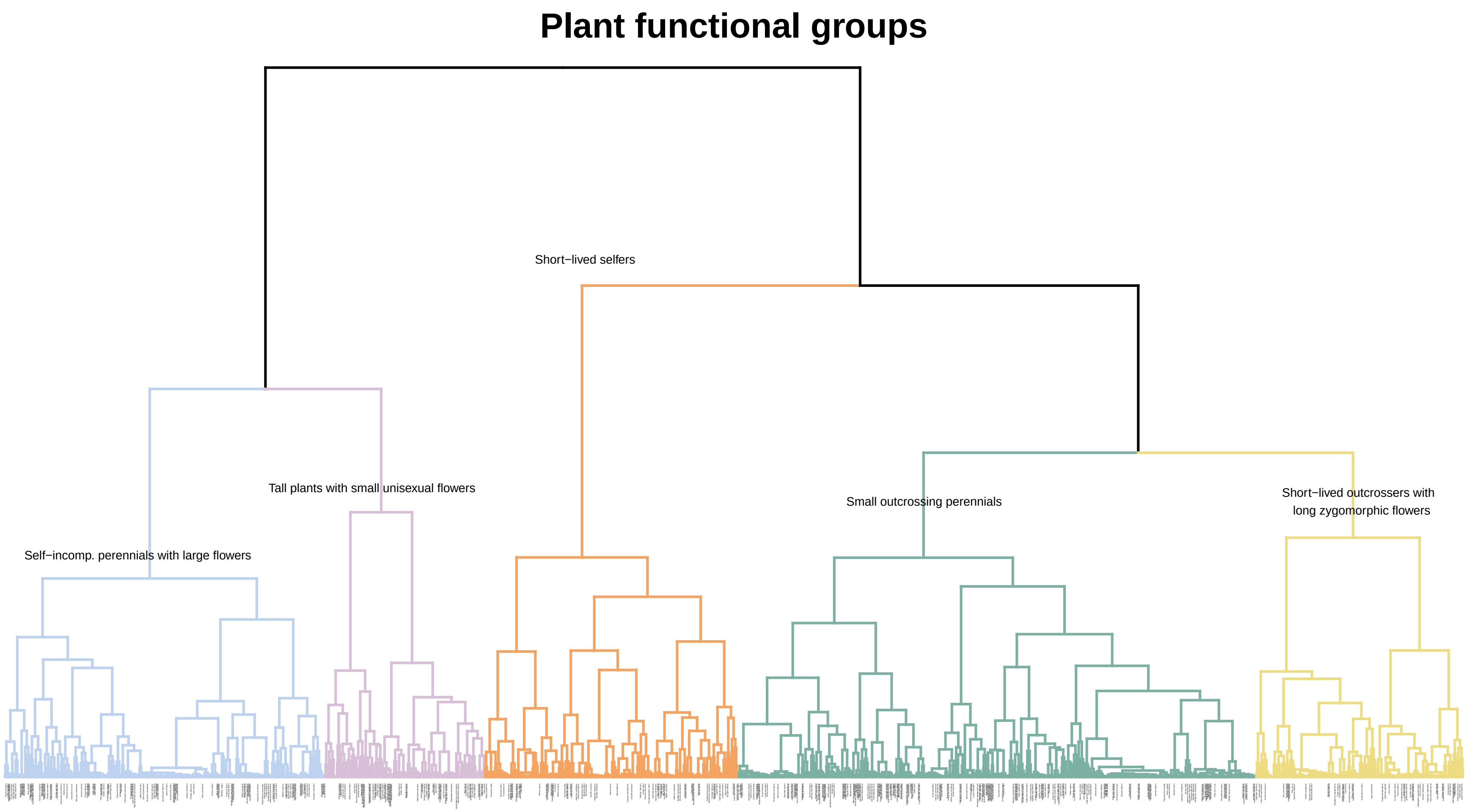


**Figure S5.** Hierarchical clustering dendrogram with the branches coloured by the optimal number of clusters (5). The labels of the subgroup of species (N = 524) used in this study are coloured in black in order to show the evenness of the distribution of the species across clusters. The rest of species labels are omitted for visualization purposes (N = 982).

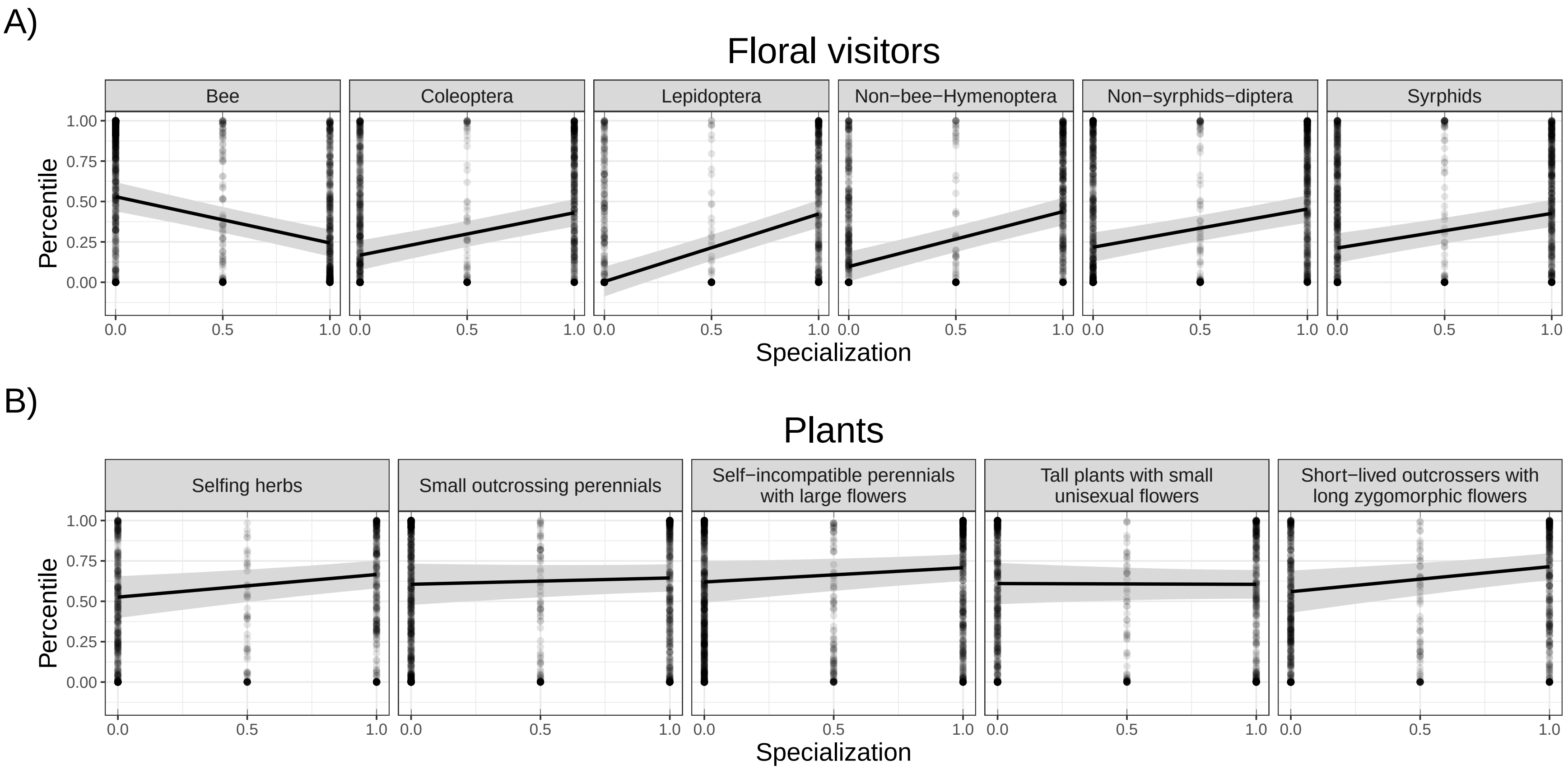


**Figure S6.** Association between the percentile (see main text) and specialization of the different plant and floral visitor groups on motif positions. Here we consider that percentiles above 0.75 indicate over-representation and percentiles below 0.25 under-representation in the comparison between empirical and simulated networks.

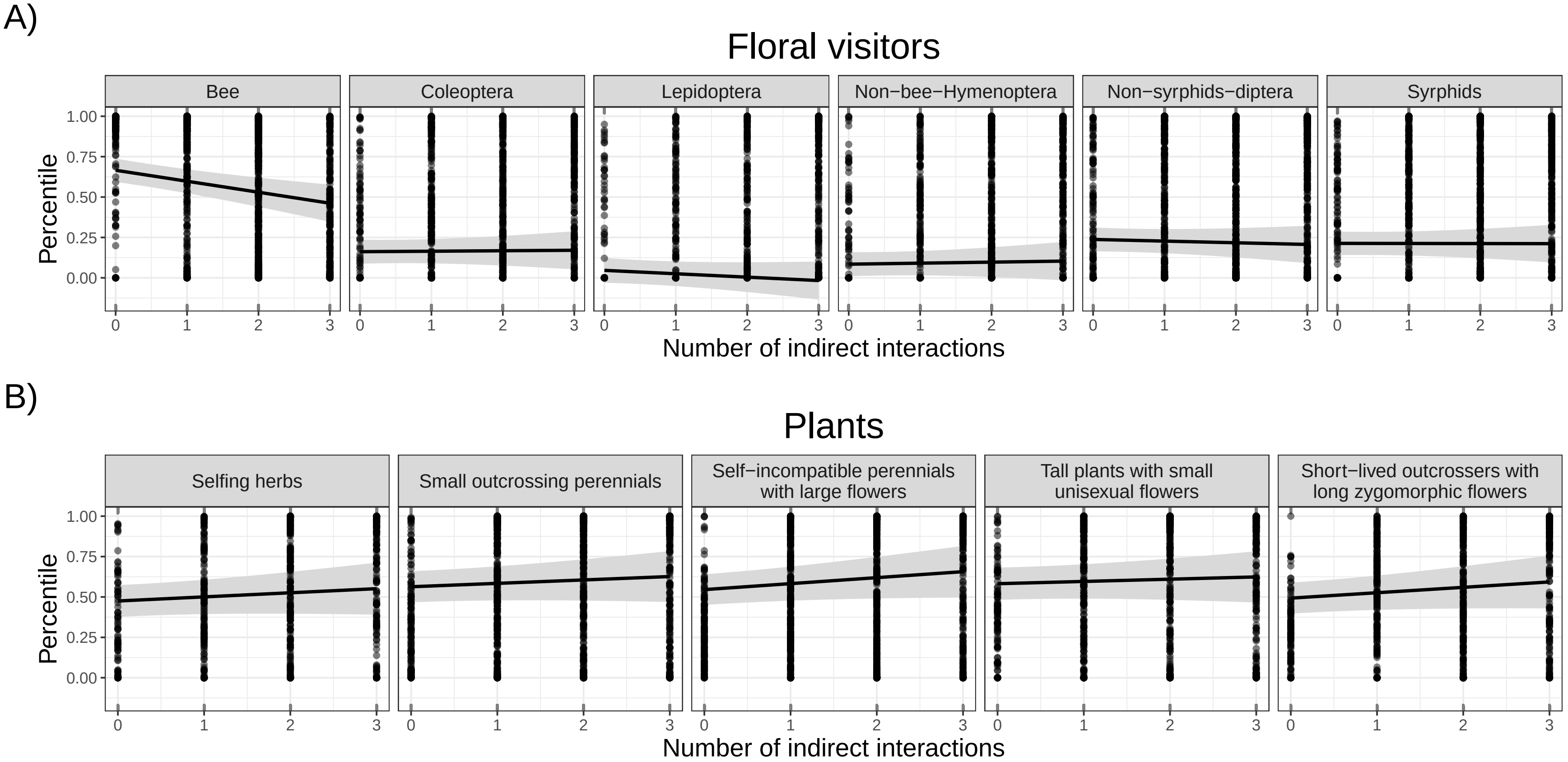


**Figure S7.** Association between the percentile (see main text) and number of indirect interactions of the different plant and floral visitor groups on motif positions. Here we consider that percentiles above 0.75 indicate over-representation and percentiles below 0.25 under-representation in the comparison between empirical and simulated networks.

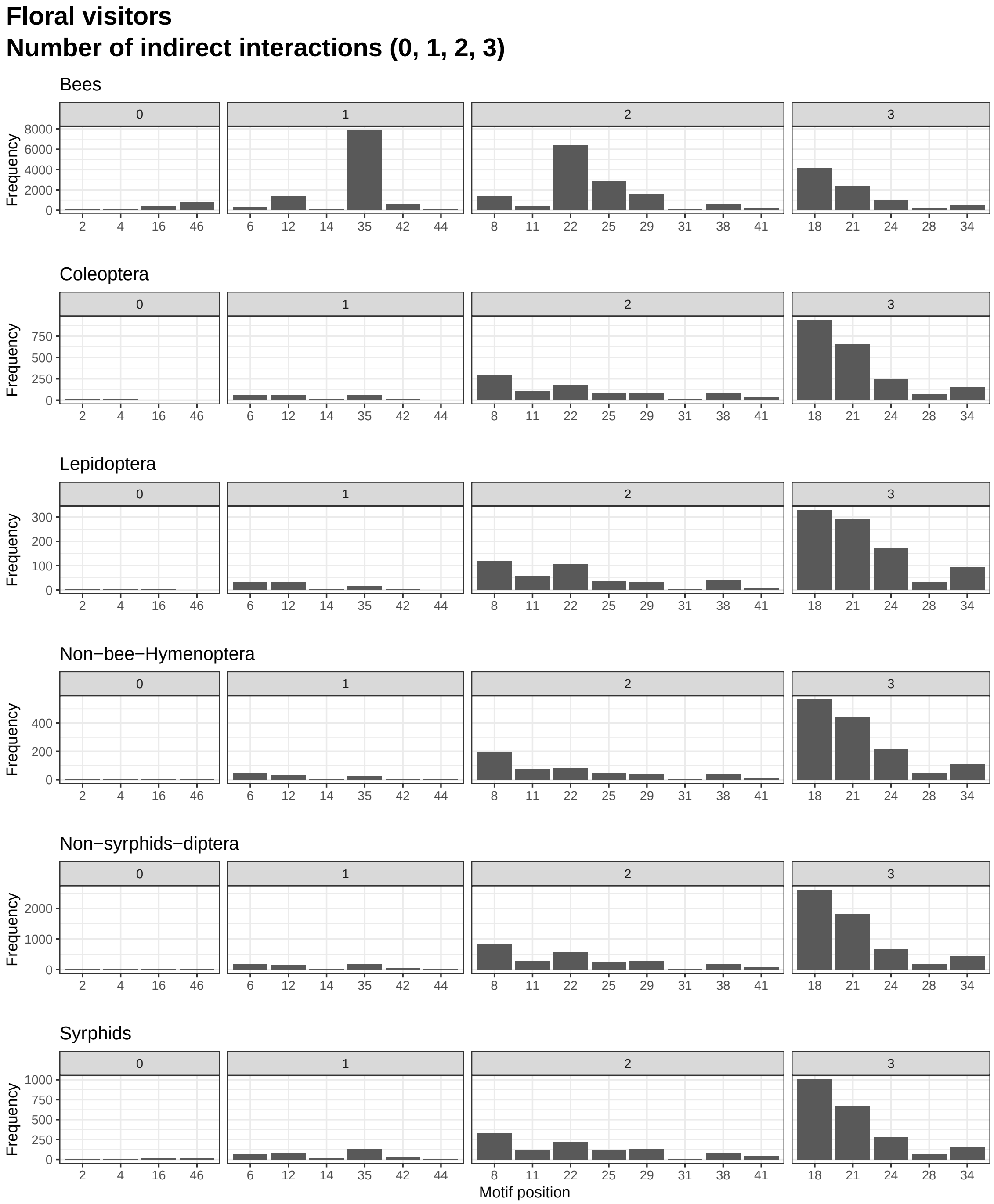


**Figure S8.** Frequency of the different floral visitor groups on the different motif positions aggregated by the number of indirect interactions for the different motif positions.

#### Plants
#### Number of indirect interactions (0, 1, 2, 3)

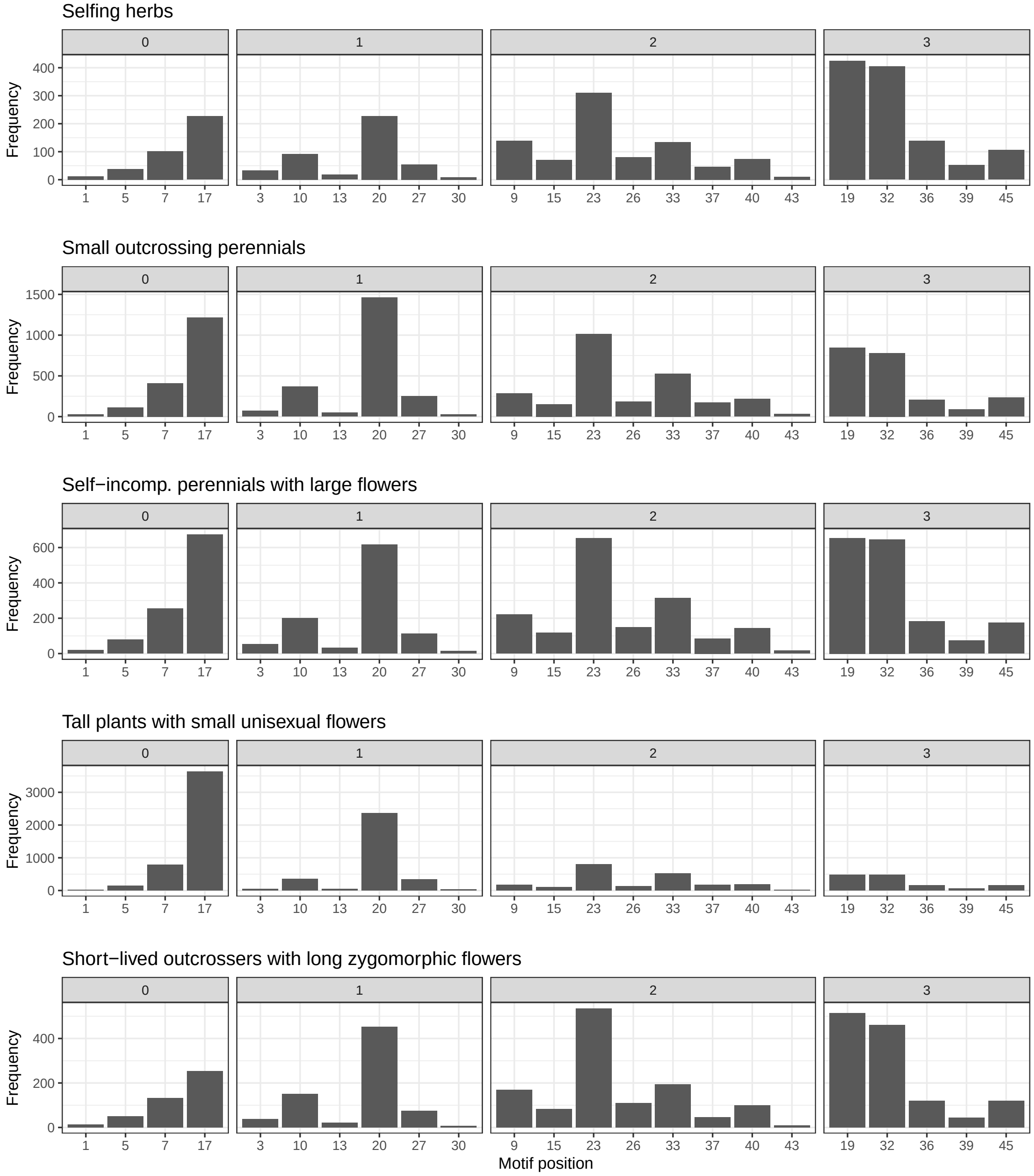


**Figure S9.** Frequency of the different plant groups on the different motif positions aggregated by the number of indirect interactions for the different motif positions.
